# TGF-β induced CXCL13 in CD8+ T cells is associated with tertiary lymphoid structures in cancer

**DOI:** 10.1101/303834

**Authors:** HH Workel, JM Lubbers, R Arnold, T Prins, P van der Vlies, K de Lange, T Bosse, I van Gool, FA Eggink, MCA Wouters, FL Komdeur, CL Creutzberg, A Kol, A Plat, M Glaire, DN Church, HW Nijman, M de Bruyn

## Abstract

Coordinated immune responses against human tumors are frequently characterized by tertiary lymphoid structures (TLS) which predict improved prognosis. The development of TLS is dependent on the chemokine CXCL13, reported to be secreted by dendritic cells and follicular helper T cells only. We report the unexpected finding that CXCL13 is also secreted by activated CD8+ T cells following stimulation by transforming growth factor beta (TGF-β). Using single cell RNA sequencing we found that expression of *CXCL13* in CD8+ T cells was restricted to the intraepithelial CD103+ population. Accordingly, CD8+ T cells activated in the presence of TGF-β simultaneously upregulated CD103 and secreted CXCL13. *CXCL13* expression was strongly correlated with neo-antigen burden and cytolytic gene signatures in bulk tumors. In line with this, TLS were abundant in neo-antigen-high, CD103+ T cell-enriched tumors. TGF-β thus appears to play a role in coordinating immune responses against human tumors through CD8-dependent CXCL13-associated formation of TLS.

## Background

Immune checkpoint inhibitors targeting programmed death ligand 1 (PDL1) or its receptor, programmed death 1 (PD-1), have elicited unprecedented long-term disease remissions in advanced and previously treatment-refractory cancers.^1–5^ Unfortunately, only a subset of patients currently benefit from treatment. Immune checkpoint inhibitors are more likely to be effective in patients with a pre-existing anti-cancer immune response; most notably a CD8+ cytotoxic T cell response against tumor neo-antigens^6^.

Responsive tumors harbor significantly more predicted neo-antigens^7,8^ and display evidence of a highly-coordinated immune response comprising T cells, dendritic cells and B cells^9^. In diseases that parallel tumor development such as chronic inflammatory conditions, this coordinated infiltration by different immune cell subsets is frequently associated with tertiary lymphoid structures (TLS) – an ectopic form of lymphoid tissue. TLS exhibit features of regular lymph nodes, including high endothelial venules, a T cell zone with mature dendritic cells (DCs) and a germinal center with follicular DCs and B cells^10^. Several studies have reported the presence of TLS in tumors, which was generally found to be associated with greater immune control of cancer growth and improved prognosis^11–14^. Furthermore, it was found for several malignancies that particularly the combination of TLS presence with high CD8+ T cell infiltration was associated to superior prognosis, whereas the presence of high CD8+ T cell infiltration alone was associated to poor or moderate prognosis^15,16^. These observations highlight the importance of a coordinated immune response, including TLS formation, in anti-cancer immunity.

To date, the molecular determinants of tumor TLS formation remain incompletely understood. Current data suggest that TLS formation results from a complex interplay between DCs, T cells, B cells and supporting stromal cells, with reciprocal signaling between these cells mediated by cytokines including chemokine [C-X-C motif] ligand 13 (CXCL13), receptor activator of nuclear factor κ B (ligand)(RANK/RANKL), lymphotoxin αβ (LΤαβ) and chemokine (C-C motif) ligand 21 (CCL21)^17,18^. A central role for CXCL13 in this process is suggested by the inability of CXCL13-knockout mice to enable homing and accumulation of B cells into lymphoid aggregates^19^ and generate functional lymphoid tissue^20,21^, and the observation that CXCL13 alone is sufficient to generate lymphoid tissue^22–24^. Nevertheless, a key outstanding question remains whether tumor-associated TLS are formed in response to the general inflammatory character of the tumor micro-environment, or rather, are induced by (neo-)antigen-specific adaptive immunity.

Here, we report on the unexpected finding that human tumor-infiltrating CTLs can produce CXCL13, linking adaptive immune activation to the formation of TLS. Notably, induction of CXCL13 in CTLs was dependent on concurrent T cell receptor (TCR)- and TGF-β receptor-signaling and was paralleled by upregulation of CD103, a marker for tissue-resident CTLs. Accordingly, the presence of CD103+ CTLs was strongly correlated to *bona fide* TLS in tumors with a high mutational load. This discovery sheds new light on how TLS could be induced by CTLs, identifying a novel role for CTLs in the orchestration of a coordinated immune response against human (neo-)antigen-rich tumors. In addition, our finding identifies CD103 and TLS as potential new biomarkers for immune checkpoint inhibitors in epithelial malignancies.

## Results

### Tertiary lymphoid structures are associated with high mutational load and increased cytotoxic T cell responses

Conflicting reports exist on the link between neo-antigen load and the presence of tertiary lymphoid structures (TLS)^16,25^. Therefore, we first assessed whether tumors with a high mutational load were enriched for TLS-associated genes in mRNA sequencing data from The Cancer Genome Atlas (TCGA). Across malignancies, a previously reported TLS gene signature^13^ was enriched in tumors with high numbers of mutations (Figure 1A). Interestingly, differences in expression of TLS genes was also observed in uterine cancer according to their molecular classification. In brief, four distinct molecular subtypes can be distinguished in uterine cancer: microsatellite stable (MSS), microsatellite unstable (MSI), Polymerase Epsilon Exonuclease Domain Mutated (*POLE*-EDM) tumors and p53-mutant tumors. We have previously demonstrated an increased number of mutations, predicted neo-antigens and cytotoxic T lymphocytes (CTLs) in *POLE*-EDM and MSI tumors compared to MSS^26^. In line with the above, MSS tumors mostly lacked TLS-related genes, while MSI and *POLE*-EDM tumors were highly enriched for TLS genes (Figure 1B). To confirm these findings, we further analyzed an independent cohort of MSS, MSI and *POLE*-EDM tumors for the presence of TLS by immunohistochemistry. In line with the TCGA data, only 48% (20/42) of MSS tumors were found to have TLS, whereas 74% (28/38) of MSI and 92% (33/36) of *POLE*-EDM tumors contained TLS (Exemplified in Figure 2A). Moreover, quantification per tumor revealed a significant increase in the number of TLS when comparing MSS to *POLE*-EDM and MSI to *POLE*-EDM tumors (Figure 2B, p<0.001 and p<0.01, respectively). To confirm that the observed CD20+ structures were *bona fide* TLS, we performed multi-color immunofluorescence. The structures we observed contained all characteristics of lymphoid tissue, as determined by the presence of high endothelial venules (HEVs), germinal B cell centers and dendritic cells (DCs) surrounded by a rim of T cells (Figure 2C). In accordance with the link between TLS and mutational load, we also identified a correlation between TLS-associated genes and CTL characteristics in TCGA data (Figure 3). Taken together, our findings suggest a link between mutational load, corresponding CTL responses and formation of TLS in human cancer.

**Figure 1.**
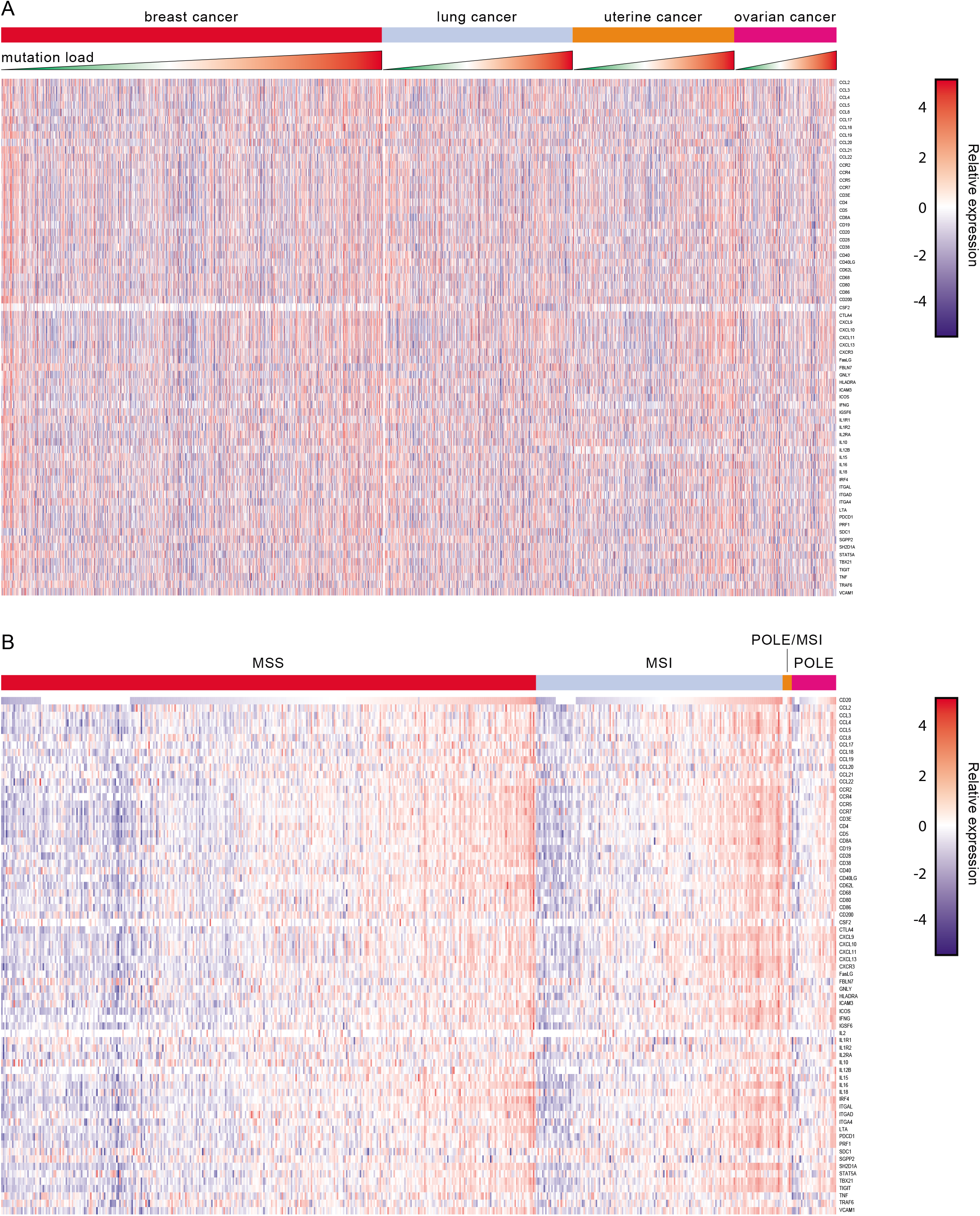
Tertiary lymphoid structures are associated with mutational load in human tumors. **A)** Heatmap of relative gene expression of tertiary lymphoid structures-associated genes, cytotoxic T cell-, CD8+ T cell- and CD4+ follicular helper T cell-related genes in TCGA data of ovarian, uterine, lung and breast cancer. Tumors are ranked from lowest to highest number of mutations from left to right. **B)** Heatmap of relative gene expression of tertiary lymphoid structures-associated genes, cytotoxic T cell-, CD8+ T cell- and CD4+ follicular helper T cell-related genes in TCGA data of uterine cancer (UCEC). Tumors are ranked according to molecular subtype.

**Figure 2.**
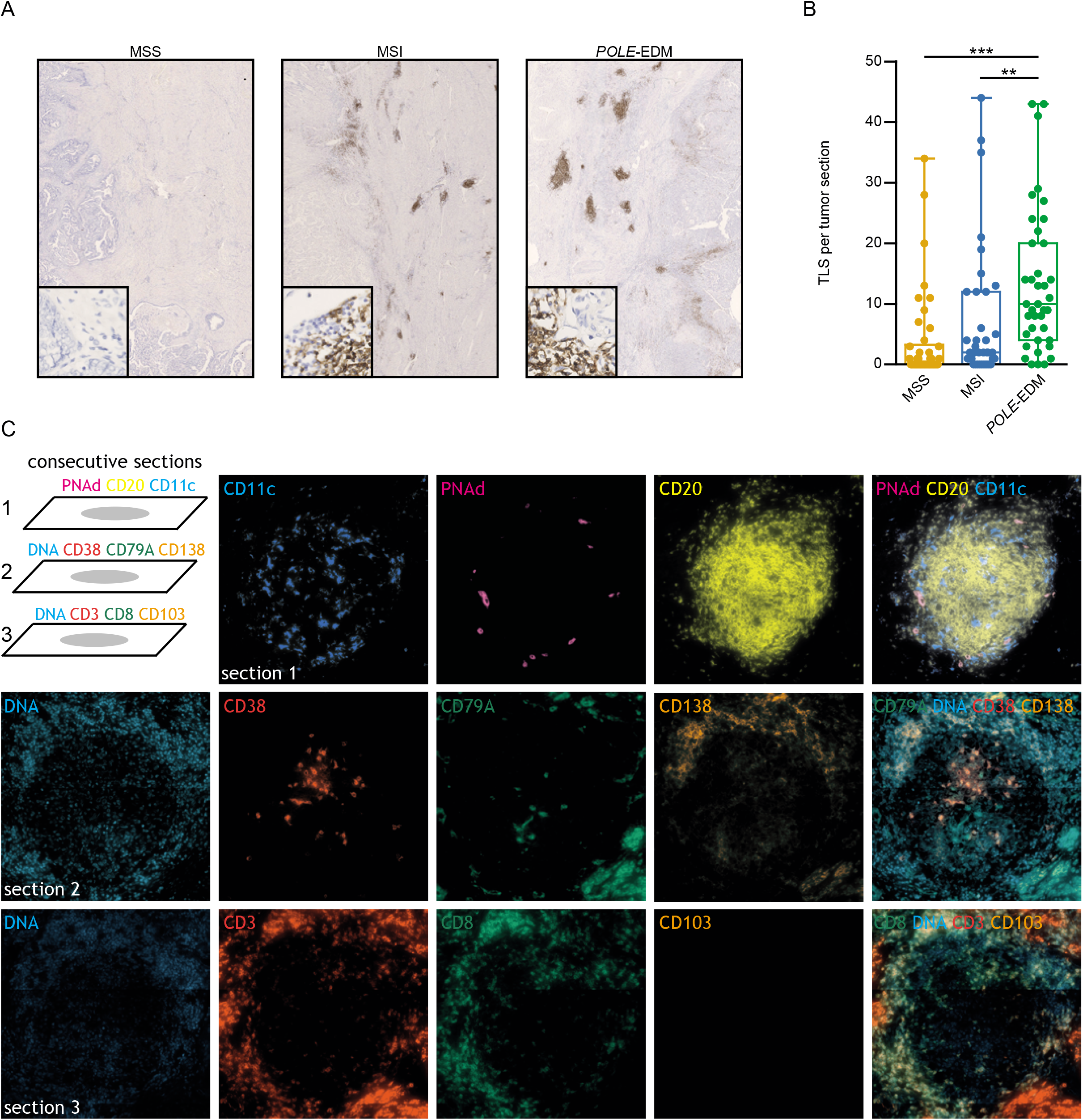
Tertiary lymphoid structures are enriched in genomically unstable uterine tumors. **A)** Representative images of immunohistochemistry for CD20 in molecular subgroups of uterine cancer, namely microsatellite stable (MSS), microsatellite unstable (MSI) and Polymerase Epsilon Exonuclease Domain Mutated (*POLE*-EDM). **B)** Quantification of tertiary lymphoid structures (TLS) in MSS, MSI and *POLE*-EDM endometrial tumors (***p<0.001 and **p<0.01). P values were calculated with a Kruskal-Wallis comparison with a post-hoc Dunn’s test. Error bars represent median±range. **C)** Multi-color immunofluorescence of TLS in a *POLE*-EDM tumor stained with three panels of tertiary lymphoid structure markers. Peripheral node addressin (PNAd, marker for high-endothelial venule), CD20 (B cells) and CD11c (dendritic cells) in panel 1 (top row), B cell markers CD38, CD79a and CD138 in panel 2 (middle row) and T cell markers CD3, CD8 and CD103 in panel 3 (bottom row).

**Figure 3.**
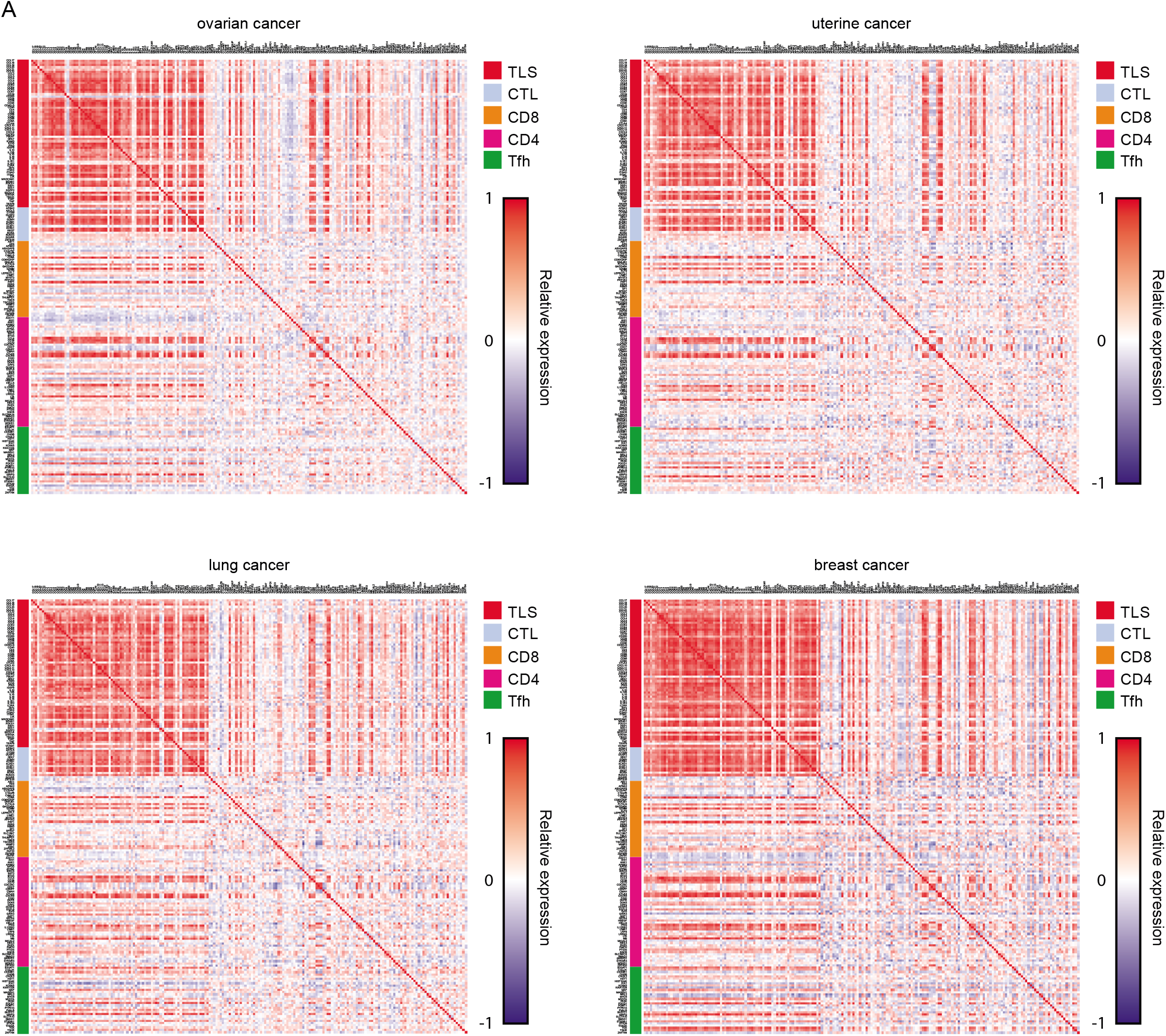
Tertiary lymphoid structure genesets are enriched in genomically unstable uterine tumors in TCGA. Spearman correlation plots of TLS, CTL, CD8, CD4 and TFH gene signatures of ovarian, uterine, lung and breast cancer log2+1 transformed mRNA sequencing data from The Cancer Genome Atlas. Relative gene expression is depicted.

### Epithelial localization of tumor-infiltrating CD8+ T cells is associated with an activated and exhausted transcriptional signature

Based on the observation that TLS-gene expression was higher in tumors with high mutational load and high CD8+ T cell infiltration, we hypothesized that tumor-reactive CTLs might be involved in the formation of TLS in cancer. To address this hypothesis, we performed mRNA sequencing on single- and 20-cell pools of CD8+ T cells isolated from human tumors. As the association between tumor mutational load, CTLs and TLS was uniform across malignancies, we chose ovarian cancer as our model tumor for the sequencing because of its large tumor bulk, high number of infiltrating, neo-antigen recognizing CD8+ cells and documented presence of TLS^16,27^. As TLS are frequently found within the tumor stroma, we also distinguished stromal from intraepithelial CD8+ T cells using the αE integrin subunit (CD103). We and others have previously shown that intraepithelial, but not stromal, CD8+ CTLs express CD103^28–31^. In line with this, TLS associated T cells were negative for CD103 (Figure 2C). CTLs were defined based on a CD3+/TCRαβ+/CD8αβ+/CD56-/CD4- phenotype (Figure 4). Post hoc t-distributed stochastic neighbor embedding (t-SNE) confirmed the presence of unique CD103+ and CD103- CTL populations in these tumors that were correctly identified by manual gating during isolation (Figure 5A). Importantly, the transcriptome of CD103+ CTLs was characterized by a marked activation and exhaustion signature with significant upregulation of *GZMB* (Granzyme B), *HAVCR2* (T cell Immunoglobulin and Mucin Domain 3, TIM3), *LAG3* (Lymphocyte-Activation Gene 3), *TNFRSF18* (Glucocorticoid-Induced TNFR-related Protein, GITR), *KIR2DL4* (Killer cell Immunoglobulin-like Receptor 2DL4), *TIGIT* (T cell Immunoreceptor with Ig and ITIM Domain) and *CTLA4* (Cytotoxic T-Lymphocyte Attenuator 4) in the 20-cell pools (Figure 5B and 5C). In addition, CD103+ CTLs expressed *GNGT2* (G Protein Subunit Gamma Transducin 2), encoding a G protein gamma family member expressed in lymph nodes and spleen that is involved in GTPase activity (Figure 5B). Differential expression of many of these markers was also observed in the single cell data (Figure 5D). The expression of these markers are in line with our earlier work demonstrating that the intraepithelial CD103+ cells represent CTLs that have undergone activation and/or exhaustion^28,29^. By contrast, CD103- CTLs displayed a more quiescent phenotype with a notable high differential expression of the V-Set Domain Containing T Cell Activation Inhibitor 1 (*VTCN1*), a known suppressor of T cell function (Figure 5B). In addition, these cells differentially expressed *GAGE12H, GAGE12I*, and *GMPR2* (Guanosine Mono Phosphate Reductase 2), involved in cell energy metabolism (Figure 5B).

**Figure 4.**
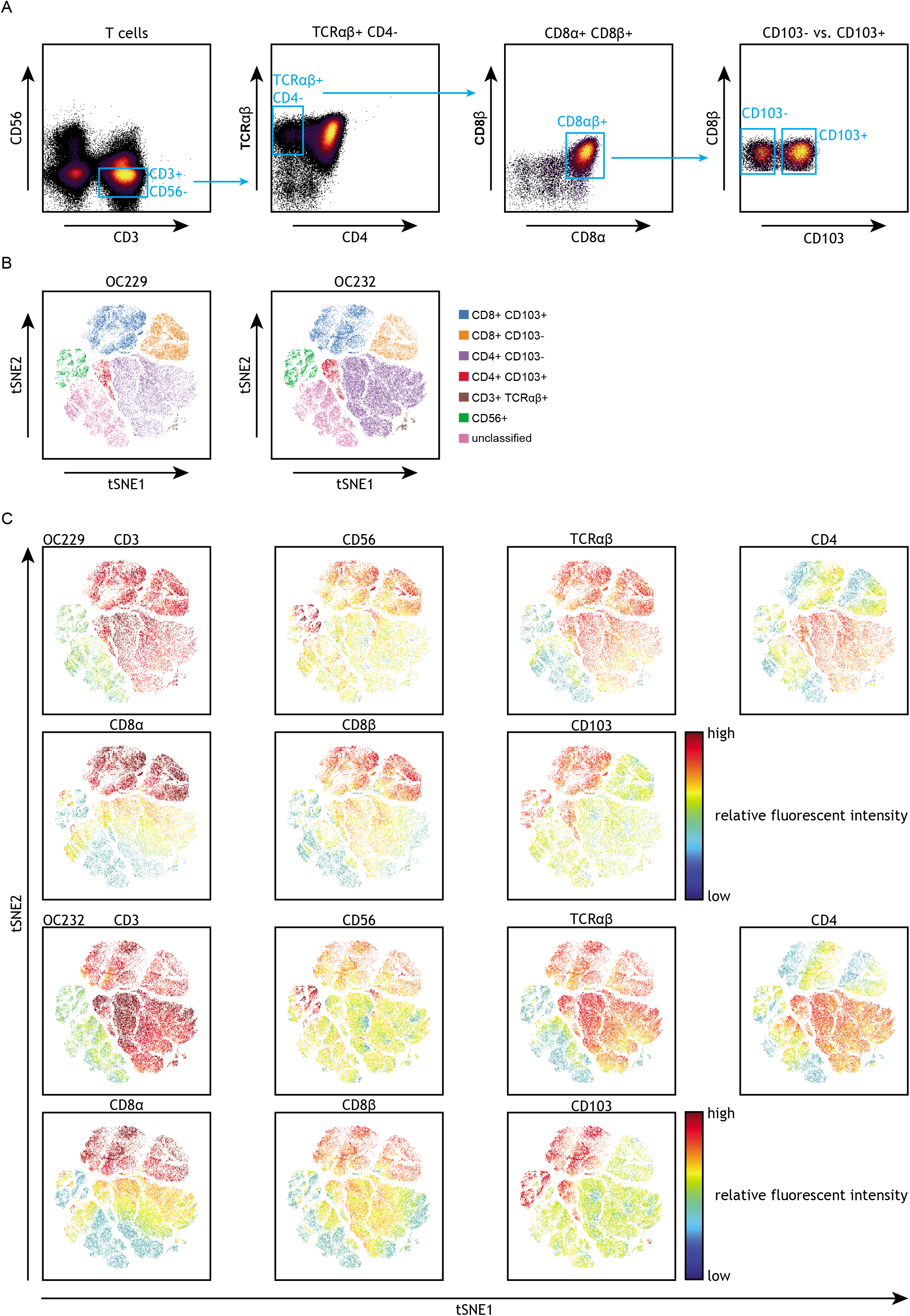
Sorting strategy for the identification of CD103+ and CD103- CD8+ T cells from primary human tumors. **A)** Gating strategy used to sort CD103+/- CD8+ T cells from human ovarian tumors. **B)** t-Distributed Stochastic Neighbour Embedding (tSNE) of flow cytometry data of individual human ovarian tumors. **C)** Relative fluorescent intensity on tSNE per flow cytometry marker, shown for individual tumors.

**Figure 5.**
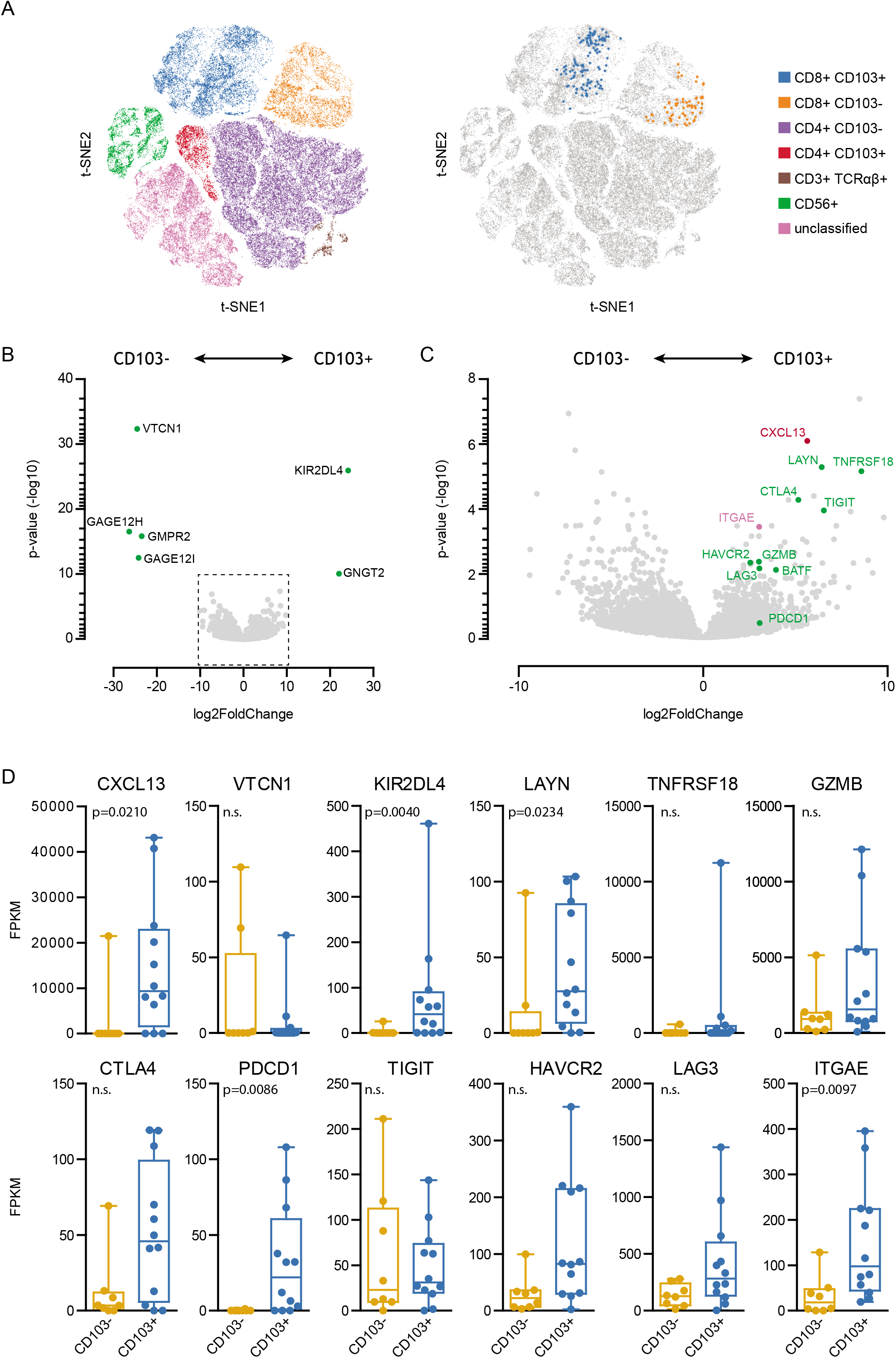
CD8^+^ CD103^+^ T cells are an exhausted T cell subtype characterized by CXCL13 expression. **A)** t-Distributed Stochastic Neighbor Embedding (tSNE) of flow cytometry data of human ovarian tumors based on flow cytometric analysis of TCRαβ CD3, CD4, CD8α, CD8β CD103 and CD56. In the right panel, the colored dots mark CD103^+^ and CD103- cells sorted for single-cell sequencing. **B)** Differential expression analysis (DESeq2) of 20-cell pool CD103^+^ versus CD103^−^ CD8+ T cell populations. Data is shown in a Volcano plot, with log2-fold change on the x-axis and the p-value on the y-axis. The highest differentially expressed (DE) genes in both the CD103^−^ and the CD103+ populations are highlighted. **C)** Inset of differentially expressed genes from panel B. DE genes involved in T cell activation and exhaustion are highlighted (green, all p-value <0.01 and a log2-fold change >2). CXCL13 is highlighted in red. As an internal control ITGAE, the gene encoding CD103, is depicted in pink. **D)** Gene-expression (Fragments Per Kilobase of transcript per Million mapped reads, FPKM) in CD103^+^ (blue) and CD103^−^ (orange) CD8^+^ single-cells. Differences were determined by a Mann-Whitney U test.

### CD103+ CTLs differentially express the B cell recruiting chemokine CXCL13

In addition to the activated and exhausted gene signature, CD103+ CTLs were also characterized by a significantly upregulated expression of the TLS-inducing chemokine [C-X-C motif] ligand 13 (*CXCL13*)(Figure 5C p< 0.0001 and Figure 5D p=0.02). This finding is of interest, as *CXCL13* is traditionally described as the prototypical CD4+ follicular helper T cell gene and since it plays a dominant role in TLS formation via recruitment of B cells and TFH through C-X-C chemokine receptor type 5 (CXCR5)^32,33^. In accordance, CXCL13 mRNA expression in uterine cancer was substantially higher in TCGA data of TLS-rich MSI and POLE-EDM tumors (Figure 1B). Since the expression of *CXCL13* in CD103+ CTLs was unexpected, we next determined whether *CXCL13* gene expression was more strongly associated with CTLs, CD8+, CD4+ or follicular helper T cell gene signatures in two gynecological malignancies, uterine and ovarian cancer. In line with our data, *CXCL13* high ovarian tumors (>median *CXCL13* expression) were strongly enriched for a CTL signature (Enrichment Score (ES) 0.93, p<0.0001). By contrast, the enrichment for total CD8+, CD4+, and TFH cell signatures was considerably lower in *CXCL13* high ovarian tumors (ES 0.72, ES 0.74, ES 0.59, respectively, all p<0.0001). Similar results were obtained in endometrial cancer (CTL ES 0.86„ CD8 ES 0.75, CD4 ES 0.75, TFH ES 0.73. In line with the observed enrichment of *CXCL13* mRNA in prognostically beneficial CD103+ CTLs, *CXCL13* gene expression was associated with a significantly improved prognosis in ovarian and uterine cancer (ovarian p=0.0188, ovarian no residual disease p=0.0394, uterine p=0.00869). As such, our data suggests CD8+ T cells may aid the formation of TLS through CXCL13.

### TGF-β primes cytotoxic CD8+ T cells to secrete CXCL13 *in vitro*

As CD8+ T cells expressed CXCL13 mRNA *in situ*, we proceeded to study whether tumor-infiltrating lymphocytes could indeed produce CXCL13 protein. CD103+ CTLs isolated from tumors of three ovarian cancer patients readily secreted CXCL13 upon *ex vivo* activation with anti-CD3/anti-CD28-conjugated beads or PMA/Ionomycin (Figure 6A). Next, we sought to define the molecular mechanism underlying the production of CXCL13. Previously, we and others have demonstrated that induction of CD103 in CD8+ T cells is dependent on concurrent T cell receptor (TCR) and Transforming Growth Factor Beta (TGF-β) receptor 1 (TGF-βR1) signaling^29,34,35^. The previously reported role of TGF-β in inducing exhaustion related genes such as PD-1 in T cells^36^ is also in line with the transcriptional profile that we obtained for CD103+ CTLs. Therefore, we hypothesized that TGF-β might prime activated peripheral blood CD8+ CTLs to secrete CXCL13. To investigate this, we activated peripheral blood CD8+ T cells from healthy donors with anti-CD3/anti-CD28-conjugated beads in the presence or absence of recombinant TGF-β1 (rTGF-β1) and measured secretion of CXCL13. Activation of CD8+ T cells alone did not induce CD103 surface-expression (Figure 6B), nor the production and secretion of CXCL13 (Figure 6C). However, in the presence of rTGF-β1, activated CD8+ T cells expressed CD103 on their surface (Figure 6B) and secreted high levels of CXCL13 (Figure 6C). Expression of CD103 and secretion of CXCL13 was inhibited by co-incubation with a TGF-βR1 inhibitor (Figure 6B and 6C). Similar results were obtained when CD8+ T cells were activated using phytohaemagglutinin (PHA) (data not shown). As interleukin 2 (IL2) inhibits the secretion of CXCL13 in follicular helper CD4+ T cells^32^, we also examined whether IL2 impacted CXCL13 secretion by CD8+ T cells. In contrast to CD4+ cells, induction of CXCL13 in CD8+ T cells was not inhibited by IL2 (Figure 6D). Next, we assessed the dose-response relationship between TGF-β and CXCL13 production and found that CXCL13 production was already significantly induced in activated CD8+ T cells at 0,1 ng/mL TGF-β and peaked at 10 ng/mL TGF-β (Figure 6E). Thus, based on our findings, we conclude that low concentrations of TGF-β are already sufficient for CXCL13 induction in CD8+ T cells. Off note, CD8+ T cells that were stimulated long-term with beads, TGFβ and IL2 maintained the ability to produce CXCL13 chemokine (not shown). To determine whether TGF-β also primes activated T cells for the secretion of other chemokines, we analyzed release of 47 chemokines from CD8+ T cells in the presence or absence of rTGF-β1 using two independent chemokine arrays (Figure 6E and 6F). As anticipated, CXCL13 was induced specifically upon activation of T cells in the presence of rTGF-β1 (Figure 6E). By contrast, no other chemokines were dependent on rTGF-β1 for their induction. Nevertheless, rTGF-β1 did appear to consistently increase the activation-dependent release of a number of chemokines. Most notably, we observed an increase in the release of chemokine (C-C motif) ligand 17 (CCL17) and chemokine (C-C motif) ligand 20 (CCL20), which are both genes that are involved in functional TLS formation (Figure 6F). In conclusion, we found that TGF-β is a specific inducer of CXCL13 production in CD8+ T cells.

**Figure 6.**
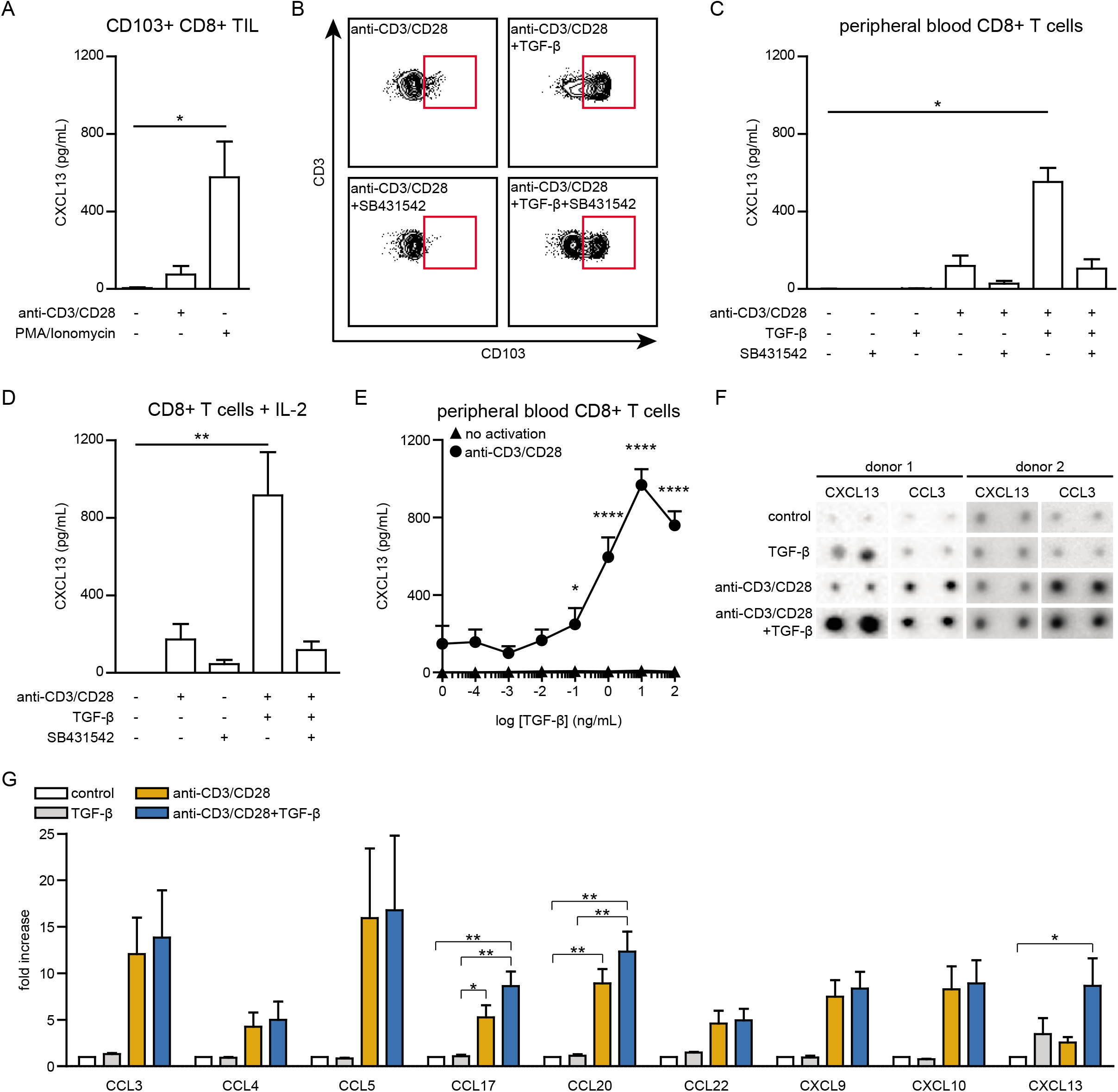
CD103^+^CXCL13^+^ T cells are induced by activation of CD8^+^ T cells in the presence of TGF-β. **A)** Tumor-infiltrating CD103^+^ CD8^+^ cells were sorted from human high-grade serous ovarian tumors (HGSOC) (n=3), cells were cultured in medium and stimulated with phorbol myristate acetate (PMA) and ionomycin, Dynabeads (αCD3/αCD28-T cell activation beads) or remained unstimulated. CXCL13 production was measured by sandwich ELISA (pg/mL)(*p<0.05). P-values were calculated using a Kruskal-Wallis comparison with a post-hoc Dunn’s test. Error bars represent mean+SEM **B)** CD8+ T cells were sorted from peripheral blood of healthy donors by negative selection magnetic activated cell sorting (n=4, analyzed in duplicate). CD8+ T cells were cultured in medium with or without Dynabeads (αCD3/αCD28-T cell activation beads), recombinant TGF-β1 and/or TGF-β1 receptor inhibitor SB431542. Expression of CD103 was assessed by flow cytometry **C)** CD8+ T cells from healthy donors were obtained as described for panel B and cultured with TGF-β or SB431532 or a combination of these. CXCL13 production was measured by sandwich ELISA (pg/mL)(n=4, analyzed in duplicate) (*p<0.05). P-values were calculated using a Kruskal-Wallis comparison with a post-hoc Dunn’s test. Error bars represent mean+SEM **D)** CD8+ T cells were cultured as described for panel B, with the addition of IL-2 in the culture medium. CXCL13 production was measured by sandwich ELISA (pg/mL) (**p<0.01). P-values for were calculated using a one-way ANOVA with a Kruskal-Wallis comparison. Error bars represent mean+SEM. **E** CD8+ T cells from healthy donors were cultured with concentrations of recombinant TGF-β1 ranging from 0 to 100 ng/mL in medium with or without Dynabeads (αCD3/αCD28-T cell activation beads) and CXCL13 production (pg/mL) was measured by sandwich ELISA (n=3, analyzed in duplicate) (****p<0.0001, *p<0.05). P-values were calculated using a two-way ANOVA followed by a post-hoc Bonferroni test. Error bars represent mean+SEM **F-G)** CD8+ T cells were sorted from healthy donors by negative selection magnetic activated cell sorting (n=3). CD8+ T cells were cultured in medium alone, with addition of TGF-β or Dynabeads, or a combination of these. Chemokine arrays were used to assess chemokine production in harvested supernatants. Representative images of chemokine array membranes (F) and densitometric analysis used to quantify chemokine production per conditions are depicted as a function of mode of activation (G) (**p<0.01, *p<0.05). P-values were calculated using a Kruskal-Wallis test with post-hoc Dunn’s test. Error bars represent mean+SD.

### Tertiary lymphoid structures are associated with the CD103+ CD8+ T cell gene signature

Based on the above, we speculated that activation of CD8+ T cell in the presence of TGF-β *in situ* would result in the induction of TLS across human epithelial tumors. As such, tumors rich in CD103+ CD8+ T cells should accumulate more TLS than tumors in which this T cell population is scarce. To assess this, we analyzed TCGA mRNA expression data of ovarian, uterine, lung and breast cancer using the CD103^+^CD8^+^ and CD103^−^CD8^+^ T cell gene signatures identified by our mRNA sequencing. The TLS gene signature was strongly correlated to CD103^+^CD8^+^ T cell genes, but not to CD103-CD8+ T cell gene signatures across all four tumor types (Figure 7). In line with our data, CXCL13-high ovarian tumors (>median CXCL13 gene expression) were strongly enriched for a CD103^+^CD8^+^ signature (Enrichment Score (ES) 0.83, p<0.0001). By contrast, there was no enrichment for CD103^−^ CD8^+^ genes in CXCL13-high ovarian tumors (ES 0.26, P=0.36). Similar results were obtained in uterine cancer (CD103^+^CD8^+^ signature ES 0.88, P<0.0001, CD103^−^CD8^+^ signature ES 0.20, P= 0.88). Taken together, our data demonstrates TGF-β1 primes CD8+ T cells to produce and secrete CXCL13, and may therefore promote the formation of TLS.

**Figure 7.**
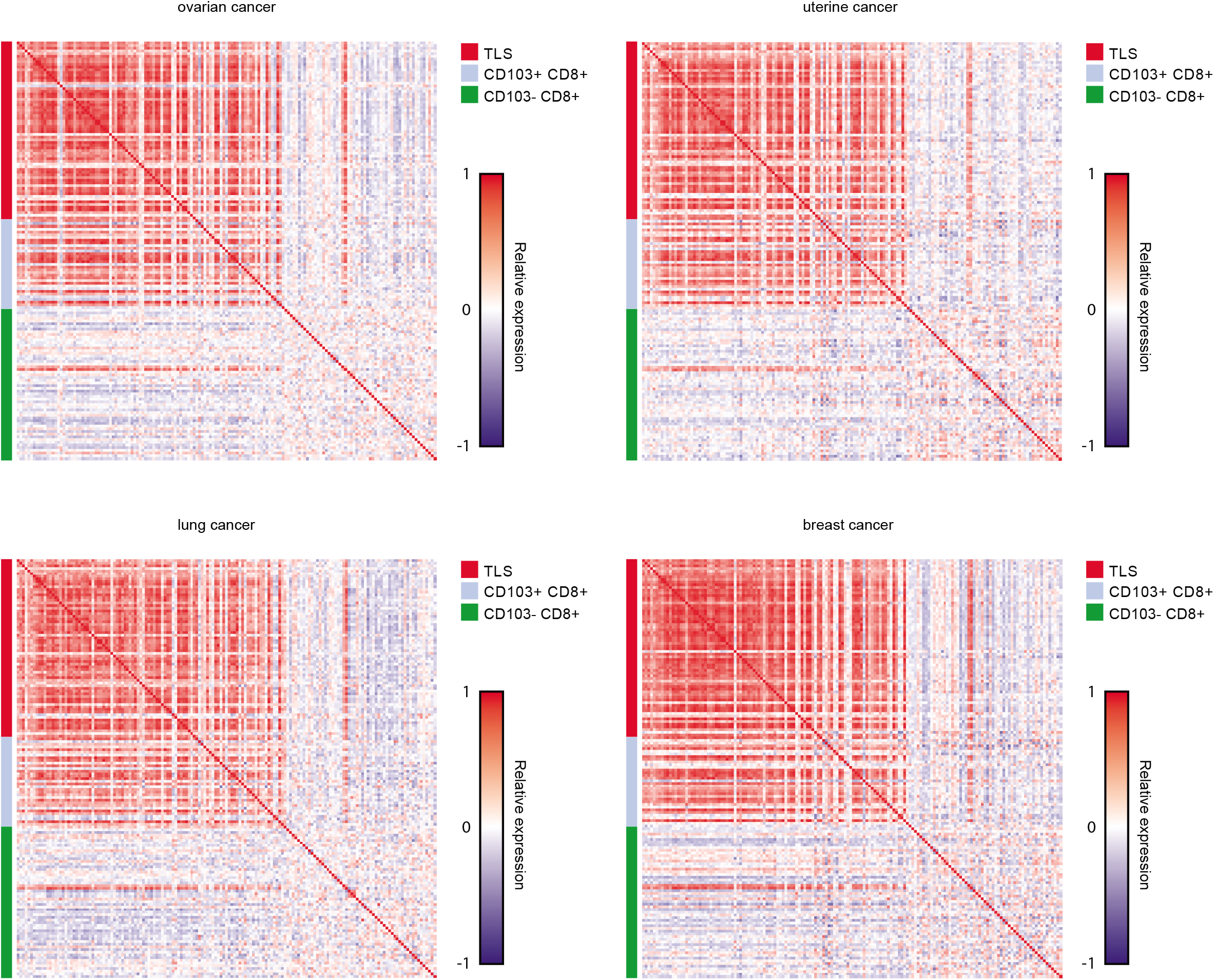
Tertiary lymphoid structures are abundant in CD103^+^CD8^+^ T cell-enriched tumors. Spearman correlation plots of TLS, CD103^+^CD8^+^ and CD103^−^CD8^+^ gene signatures of ovarian, uterine, lung and breast cancer log2+1 transformed mRNA sequencing data from The Cancer Genome Atlas. Relative gene expression is depicted.

## Discussion

In this study we report on the unexpected finding that transforming growth factor beta (TGF-β) stimulates activated CD8+ T cells to produce chemokine (C-X-C motif) ligand 13 (CXCL13), a known inducer of tertiary lymphoid structures (TLS)^22–24^. This production of CXCL13 was paralleled by the induction of CD103 on the cell surface of CD8+ cells *in vitro.* Further, CD103+ cytotoxic T lymphocytes (CTLs) isolated directly from human tumors strongly expressed *CXCL13* mRNA and secreted CXCL13 protein upon *ex vivo* reactivation. Notably, the presence of TLS gene signatures was strongly increased in highly mutated, CD103+ T cell-enriched human tumors from The Cancer Genome Atlas (TCGA). Further, the absolute number of TLS was increased in an independent cohort of neo-antigen-low, -intermediate and -high endometrial cancers. Our findings shed new light on the link between innate and adaptive immunity in general and on the link between CD8+ CTL activation and the induction of TLS in particular. Our data identify CD103 and TLS as potential biomarkers of interest for cancer immunotherapy.

The expression of *CXCL13* in CD103+ CTLs was remarkable, since CXCL13 is reported to be produced by DCs, T_FH_ and B cells only^32,33,37,38^. We have therefore carefully assessed previously published (single cell) sequencing data of exhausted, tumor-infiltrating T cells of liver cancer, lung cancer and melanoma^9,39,40^. These published data support our finding that exhausted CD8+ TIL can express *CXCL13* on the mRNA level, even though this finding was not mentioned in either of the papers. Moreover, the finding that TGF-β1, a cytokine mostly associated with immune suppression^32,41–45^, was essential for the induction of CXCL13 is intriguing. Under homeostatic conditions, TGF-β1 is abundantly present in epithelial tissue and controls the epithelial localization of resident memory immune subsets such as the intraepithelial lymphocytes in the colon^46^. In epithelial cancers, we suggest that TGF-β1 has a similar role in promoting not only recruitment, signaling and retention of CD8+ CTLs via CD103 expression^35^, but also stimulating immunity via attraction of C-X-C chemokine receptor type 5 (CXCR5)+ immune cells through CXCL13 signaling.

CXCL13 is the key molecular determinant of TLS formation^22–24^, ectopic lymphoid structures that are thought to enable efficient local priming of T cells by dendritic cells (DCs)^11,12^. Hereby, the time-consuming migration of DCs and T cells to and from secondary lymph nodes may be circumvented, augmenting local anti-tumor immunity. In line with this, characteristic components of TLS such as high endothelial venules (HEVs) and B cells, were found to be generally associated with an improved prognosis^13^. In addition, plasma B cells in the TLS are thought to enhance the antitumor response by production and subsequent accumulation of anti-tumor antibodies, potentially leading to antibody-dependent cytotoxicity and opsonization^16^. Thus, TLS may orchestrate a joint T and B cells response to improve anti-tumor immunity.

As TLS were found to be more abundant in tumors with a high mutational load, we postulated that activated CD103+ CTLs are involved in the formation of TLS in cancer via production of CXCL13. This is supported by the observations that highly mutated, CD8+ T cell-rich tumors showed higher expression of *CXCL13* and *ITGAE* (CD103) and that they presented with significantly higher numbers of TLS. In accordance, a higher degree of T cell receptor clonality within CD8+ T cells correlated with a higher number of TLS in non-small cell lung cancer^47^. These TLS may represent an ongoing immune response that was insufficient to halt tumor progression at an early time point. It would therefore be of great interest to study the induction and formation of TLS in developing cancer lesions and to determine whether CD8 infiltration precedes TLS formation.

In line with previous work^28,29^, CD103+ CTLs from human tumors were also characterized by a marked activation and exhaustion-related gene expression signature with differential expression of granzymes and well-known immune checkpoint molecules, such as cytotoxic T-lymphocyte-associated protein 4 (*CTLA4*). In addition, CD103+ CTLs expressed a host of additional immune checkpoint genes currently under clinical investigation, such as T cell immunoglobulin and mucin-domain containing-3 (*TIM3*), lymphocyte-activation protein 3 (*LAG3*) and T-cell immunoglobulin and ITIM domain (*TIGIT*). As such, our findings also have implications for clinical immunotherapy.

Indeed, tissue-resident CD103+ CTLs were recently found to be significantly expanded upon treatment with Nivolumab and Pembrolizumab (anti-PD-1) in tumor specimens of advanced stage metastatic melanoma patients^48^. Accordingly, a paper by Riaz et al. demonstrated that tumors from patients who responded to Nivolumab treatment differentially expressed genes such as *CTLA4, TIM3, LAG3, PDCD1*, Granzyme B (*GZMB*), tumor necrosis factor receptor superfamily member 9 (*TNFRSF9*) and *CXCL13*, all genes overexpressed in CD103+ vs. CD103- CTLs^9^. Notably, pre-treatment, but not on-treatment *CXCL13* was differentially expressed in responders vs. non-responders in this study^9^. This may be explained by the low basal CXCL13 secretion we observed in the exhausted, CD103+ CTLs freshly isolated from untreated human tumors. In their exhausted state, CTLs might accumulate mRNA encoding several key effector molecules, that is translated only upon reactivation by e.g. immune checkpoint blockade (ICB). In line with this, Riaz et al. observed a marked increase in the number of B cell-related genes on treatment in responding patients^9^, perhaps hinting at the formation of TLS in these patients upon ICB-mediated release of CXCL13. This hypothesis is supported by the recently published increase in serum CXCL13 levels and concomitant depletion of CXCR5+ B cells from the circulation in patients treated with anti-CTLA-4 and/or anti-PD-1 antibodies^49^. Our data therefore suggest that ICB is of particular interest for patients with a high CD103+ CXCL13+ CTL infiltration pre-treatment across malignancies.

Several novel combination immunotherapy regimes that promote CTL infiltration and TLS formation may also function via CTL-dependent production of CXCL13. For instance, combined therapy with anti-angiogenic and immunotherapeutic agents in mice stimulated the transformation of tumor blood vessels into intratumoral HEVs, which subsequently enhanced the infiltration and activation of CTLs and the destruction of tumor cells^50,51^. These CTLs formed structures around the HEVs that closely resembled TLS^50,51^. One of these studies found that induction of TLS was dependent on both CD8+ T cells and macrophages^50^. However, the exact intratumoral mechanism of action remained unclarified. Since macrophages produce TGF-β in a chronically inflamed environment^32^, we hypothesize that the macrophages in these studies may have generated a TGF-β enriched environment, thus leading to the production of CXCL13 chemokine by activated T cells and subsequently to the formation of lymphoid structures. TLS may therefore reflect an ongoing CD8+ T cell response in cancer. As such, TLS may be used as a biomarker to predict response to immune checkpoint blockade. In addition, these structures may be used as a general biomarker for response to immunotherapy, since TLS were found to mark pancreatic cancer patients who responded to therapeutic vaccination^52^.

Taken together, we demonstrate that TGF-β1 induces co-expression of CXCL13 and CD103 in CD8+ T cells, linking CD8+ T cell activation to TLS formation. Our findings therefore provide a new perspective on how (neo-)antigens can promote the formation of TLS in human tumors. Accordingly, TLS and/or CD103+ cells should be considered as a potential novel predictive or response biomarker for immune checkpoint blockade therapy.

## Materials and Methods

### Patients

Tumor tissue from four patients with stage IIIC high-grade serous ovarian cancer was collected during primary cytoreductive surgery, prior to chemotherapy, and from one patient with stage IV high-grade serous ovarian cancer during interval debulking upon three cycles of chemotherapy. Selection of uterine cancer (UC) patients was described previously^53^. Briefly, UC tissue was obtained from patients involved in the PORTEC-1 and PORTEC-2 studies (n=57) and the UC series (n=67) rom Leiden University Medical Center (LUMC) and UC series (n=26) from the University Medical Center Groningen^26^. Tumor material from 119 patients was available for analysis. Mutations in the exonuclease domain of polymerase epsilon (*POLE*-EDM) and microsatellite instability status were known from previous studies (Van Gool, Church). Of the tumors available for this study, 42 tumors were *POLE* wild-type, microsatellite stable (MSS), 38 were *POLE* wild-type, microsatellite unstable (MSI) and 39 were *POLE*-EDM. *POLE*-EDM statuses did not co-occur with microsatellite instability. All cases were of endometrioid histology (EEC) and the number of low grade and high-grade tumors was spread equally over the three molecular groups. Ethical approval for tumor molecular analysis was granted at LUMC, UMCG and by Oxfordshire Research Ethics Committee B (Approval No. 05\Q1605\66).

### Analysis of TCGA mRNA sequencing data

RSEM normalized mRNAseq data and clinical data from uterine corpus endometrial carcinoma (UCEC), ovarian cancer (OV), breast cancer (BRCA) and lung adenocarcinoma (LUAD) were downloaded from firebrowse.org on 13-03-2017 (UCEC) and 14-07-2017(OV, BRCA, LUAD). RSEM mRNA sequencing expression data were log2 +1 transformed and genes with zero reads in all samples were removed. *POLE*-EDM, MSI and MSS cases were identified in the endometrial cancer data; mononucleotide and dinucleotide marker panel analysis status was provided by The Cancer Genome Atlas (TCGA) and mutations in the exonuclease domain of *POLE* were determined previously^54^. Heatmaps were constructed in R (version 3.3.1) with packages gplots and ggplots. The javaGSEA Desktop Application was downloaded from http://software.broadinstitute.org/gsea/index.jsp. TCGA uterine corpus endometrial cancer and ovarian cancer log2+1 transformed data and phenotype data were converted to a suitable format and entered in the GSEA desktop application. Several gene-sets were used to determine enrichment for cytotoxic T lymphocytes (CTLs), CD8+ T cells, CD4 helper cells or tertiary lymphoid structures(TLS)^13,55^, as well as the CD8+CD103+ signatures derived from the sequencing data. Gene-set enrichment for TLS genes was determined for *POLE*-EDM versus MSS in uterine corpus endometrial cancer and gene-set enrichments for CTLs, CD8, CD4, TFH and CD8+CD103+ genes were determined for CXCL13hi versus CXCL13lo (based on median gene expression) tumors in all cancers analyzed.

Spearman correlations between the TLS signature, the CD8+CD103+ signature and the CD8+CD103-signature were visualized in correlation plots using the Corrplot package (Version 0.77) in R. Differences in survival were evaluated with a logrank test within the Survival package (Version 2.41-3) in R. All analyses were performed in R (version 3.4.0), with exception of the construction of the heatmap in Figure 1, which was made in R version 3.3.1.

### Immunohistochemistry

Formalin-fixed, paraffin-embedded (FFPE) slides were de-paraffinized and rehydrated in graded ethanol. Antigen retrieval was initiated with a preheated 10mM citrate buffer (pH6) and endogenous peroxidase activity was blocked by submerging sections in a 0.45% Hydrogen peroxide solution. Slides were incubated overnight with 0.63 mg/L of anti-CD20 antibody (Dako, Glostrup, Denmark) at 4°C. Subsequently, slides were incubated with a peroxidase-labeled polymer for 30 minutes (Envision+ anti-mouse Dako, Carpinteria, USA). Signal was visualized with 3,3’diaminobenzidin (DAB) solution and slides were counterstained with haematoxylin. Appropriate washing steps with PBS were performed in-between incubation steps. Sections were embedded in Eukitt mounting medium (Sigma Aldrich, Steinheim, Germany) and slides were scanned on a Hamamatsu digital slide scanner (Hamamatsu photonics, Hamamatsu, Japan). The number of CD20+ (dense) follicles in each slide was quantified in NDPview2 software by two independent observers who were blinded to clinicopathological data. Immunohistochemistry for CD8 was performed previously in this cohort^26^.

### Multi-color immunofluorescence

FFPE slide preparation and antigen retrieval were performed as described above. Next, slides were incubated overnight at 4°C with primary antibody and subsequently incubated with the appropriate secondary antibody for 45 minutes at room temperature. Specific signal was amplified using the TSA Cyanine 5 (Cy5) detection kit (Perkin Elmer, NEL705A001KT, Boston, USA) or the TSA Cyanine 3 (Cy3) and Fluorescein detection kit (Perkin Elmer, 753001KT, Waltham, USA), according to manufacturer’s protocols. To allow multiple amplifications on the same slide, primary HRP labels were destroyed between incubations by washing with 0.01 M hydrochloric acid for 10 minutes. Appropriate washing steps with PBS-0,05%Tween20 were performed during the procedure. Finally, slides were embedded in Prolong Diamond anti-fade mounting medium with or without DAPI (Invitrogen/Thermo Fisher Scientific, P36962 and P36961, Oregon, USA) and scanned using the TissueFAXS microscope (TissueGnostics, Vienna, Austria). Overlay images were produced using Adobe Photoshop software.

### mRNA sequencing

Ovarian tumors from two patients were cut into pieces of <1 mm3 and put in a culture flask with digestion medium, consisting of RPMI (Gibco, Paisley, UK), 10% Fetal Bovine Serum (FBS, Gibco, Paisley, UK), 1 mg/ml collagenase type IV (Gibco, Grand Island, USA) and 12.6 mL/L Pulmozyme (Roche, Woerden, the Netherlands) for overnight digestion at room temperature. After digestion, the suspension was washed with PBS and strained through a 70 μm filter. Cells were centrifuged over a Ficoll-Paque gradient (GE Healthcare Bio-Sciences AB, Uppsala, Sweden), suspended in FBS with 10% dimethylsulfoxide and stored in liquid nitrogen until further use. Prior to sequencing, tumor digests were thawed on ice, washed with AIM-V medium (Gibco, Paisley, UK) with 5% pooled human serum (PHS, One Lambda, USA) and centrifuged at 1000g. Pellets were resuspended in AIM-V with 5% PHS and cells were incubated with CD3-BV421, CD4-PerCP-Cy5.5, CD8α-APCeFluor780, CD8β-PEcy7 TCRαβ-APC, CD103-FITC and CD56-PE antibodies at 4°C for 45 minutes. After gating for CD3+CD4-CD8αβ+TCRαβ+CD56- cells, CD103- and CD103+ single cells were sorted on a Beckman Coulter Astrios directly into lysis buffer (0.2 % Triton X-100 and Recombinant RNase inhibitor (Westburg-Clontech) in 96-well PCR plates. Each well contained a unique indexed Oligo dT primer, enabling identification of individual cells after pooled RNA sequencing. In addition to single cells wells, small bulk population of 20 cells were sorted per microplate well. Per patient, 40 single CD8^+^ T cells (20 wells CD103^+^, 20 wells CD103^−^) and 20 small bulk 20-cell populations (10 wells CD103^+^, 10 wells CD103^−^) were sorted. After lysis of the cells, the transcriptomes were amplified by a modified SMART-Seq2 protocol using SmartScribe reverse transcriptase (Westburg-Clontech, CL639537), based on a previously published protocol (Picelli et al.^REF^). Sequencing libraries were prepared using the Illumina Nextera XT DNA sample preparation kit. Presence and size distribution of the obtained PCR product were checked on a PerkinElmer LabChip GX high-sensitivity DNA chip. A super pool was created by equimolar pooling of the Nextera products and the samples were sequenced on Illumina NextSeq500 2500 using 50bp paired-end reads, one read for the mRNA transcript and the other for the cell-barcode. The obtained RNA sequencing data were demultiplexed into individual FASTQ files. The obtained single-end reads were aligned to human reference genome 37 (GRCh37, top-level built), using STAR (version 2.5.2). We then used RNA-SeQC (version 1.1.8) to assess the quality of each sample and all cells that did not meet one of the following criteria were removed: <10000 transcripts detected, <500000 uniquely mapped reads, <1000 genes detected, a mapping rate of <0.5, an expression profiling efficiency of <0.4 or an exonic rate of <0.5. Differential expression was analyzed to obtain insight into the differences between CD103^+^ and CD103^−^ CD8^+^ T cells from the 20-cell populations with DESeq2 (version 1.16.1). For this analysis, expression values for each sample have been obtained using RSEM (version1.3.0, with Bowtie 2, version 2.2.5, non-stranded and with the single-cell prior activated to account for drop-out genes in both, bulk and single cells) and have been computed for the Gencode 19 transcriptome annotation for GRCh37 (reference index built with – polyA activated).

Genes with a Benjamini-Hochberg adjusted p-value of <0.05 were selected for further analysis. Differentially expressed genes were visualized in a Volcano plot (DESeq2, version 1.16.1).

### ELISA

Tumor-infiltrating lymphocytes from three high-grade serous ovarian cancer digests were stained and sorted as described for mRNA sequencing. The numbers of sorted T cells for the three patients were 163×10^3^, 216×10^3^ and 154×10^3^ for CD4+ cells, 82×10^3^, 38×10^3^ and 83×10^3^ for CD8+CD103- and 207×10^3^, 120×10^3^ and146×10^3^ for CD8+CD103+ T cells. Sorted T cells remained unstimulated or were activated, either with phorbol myristate acetate (PMA) and ionomycin (500× dilution, Invitrogen, 00-4970-93 Carlsbad USA) or with Dynabeads^®^ (2μL/1×10^5^ cells, T-activator CD3/CD28 beads, 11131D, Gibco, Oslo, Norway and Vilnius, Lithuania). In addition, peripheral blood CD8+ T cells were isolated from blood of four healthy volunteers by a Ficoll-Paque gradient followed by magnetic activated cell sorting with a CD8 T cell negative selection kit (Affymetrix, San Diego, USA). Peripheral blood CD8+ T cells were incubated in AIM-V medium, with or without Dynabeads^®^ (2μL/1×10^5^ cells) for activation, recombinant TFG-β1 (rTGF-β1, 100 ng/mL, Peprotech, USA), TGF-β1 receptor inhibitor (10μM, SB431542, Sigma Aldrich/Merck, Saint Louis, USA) or a combination of these. Similar experiments were performed with the addition of IL2 (100 IU/mL. Novartis Pharmaceuticals, UK). For the dose-response curve, peripheral blood CD8+ T cells from three healthy donors were incubated with or without Dynabeads^®^ (2μL/1×10^5^ cells) for activation and with recombinant TFG-β1 at doses ranging from 0 to 100 ng/mL (rTGF-β1, Peprotech, USA). All cells were cultured in AIM-V medium with 5% pooled human serum in 96-well plates containing 1×10^5^ cells per condition. After 7 days, plates were centrifuged and supernatant was collected for ELISA. CXCL13 sandwich ELISA experiments were performed according to manufacturer’s protocol (Human CXCL13/BLC/BCA-1 DuoSet ELISA DY801, R&D Abingdon, UK or, for the dose-response curve, Minneapolis, USA). In brief, plates were coated with a capture antibody, followed by incubation with cell supernatant. Binding of CXCL13 was detected using secondary antibody, streptavidin-HRP and TMB 1-Component Microwell Peroxidase Substrate (SureBlue, KPL/SeraCare, Milford, USA) Substrate conversion was stopped after 20 minutes with 0.01M Hydrogen Chloride. Plates were washed with PBS-0.05%Tween20 in-between incubations. OD values were obtained using a micro plate reader set to 450 nm (BioRad iMark™ Microplate reader). AIM-V medium was used as a negative control.

### Chemokine arrays

CD8+ T cells were isolated from from blood of three healthy donors as described for ELISA. Per condition, 5×10^5^ cells were cultured in AIM-V medium with 5% PHS in a 24-well plate. Cells were either incubated for 7 days in medium alone, with rTGF-β1 (100 ng/mL, Peprotech, USA), with Dynabeads^®^ (2μL/1×10^5^ cells, T-activator CD3/CD28 beads, 11131D, Gibco, Oslo, Norway and Vilnius, Lithuania) or with both rTGF-β1 and Dynabeads^®^. Samples were centrifuged and supernatants were collected to analyze production of chemokines on chemokine arrays, according to manufacturer’s instructions (31 chemokines using the Proteome Profiler Human Chemokine Array Kit, ARY017, R&D, Abingdon, UK, and 38 chemokines using the Human Chemokine Antibody Array - Membrane, ab169812, Abcam, Huissen, the Netherlands). In brief, chemokine receptor-coated membranes were incubated with supernatant over night at 4°C. Next, captured proteins were visualized using chemiluminescent detection reagents. Appropriate washing steps were performed in-between incubation steps. Membranes were imaged on BioRad ChemiDoc™ MP Imaging System, densitometric analysis of chemokine spots was performed using the Protein Array Analyzer plugin for Image J^56^.

### Statistical analyses

Differentially expressed genes in CD103+CD8+ versus CD103-CD8+ T cells sorted from human ovarian tumors were determined by DESeq2 for 20 cells-populations. Genes with a Benjamini Hochberg adjusted p-value of <0.05 were selected for further analysis. Differences in FPKM-values of single-cells were assessed by a Mann-Whitney U test. Differences in number of TLS on FFPE slides of molecular subgroups of EC were determined by a non-parametric Kruskal-Wallis test, followed by Dunn’s post-hoc analysis. We analyzed TCGA mRNA sequencing data and compared differences in gene expression between molecular subgroups of EC with a non-parametric Kruskal-Wallis test and a post-hoc Dunn’s test. CXCL13 production was analyzed using a Kruskal-Wallis comparison with a post-hoc Dunn’s test, or, for the dose-response curve, with a two-way ANOVA followed by a post-hoc Bonferroni test. The chemokine arrays were analyzed using a Kruskal-Wallis test with a post-hoc Dunn’s test. All statistical analyses were performed using R version 3.4.0 or GraphPad Prism (GraphPad Software Inc., CA, USA).

## Acknowledgements

The authors would like to thank Henk Moes, Geert Mesander, Johan Teunis, Joan Vos and Niels Kouprie for their technical assistance. This work was supported by Dutch Cancer Society/Alpe d’Huzes grant UMCG 2014–6719 to MB, Dutch Cancer Society Young Investigator Grant 10418 to TB, Jan Kornelis de Cock Stichting grants to FLK, KLB, FAE and HHW, Nijbakker-Morra Stichting and Studiefonds Ketel1 grants to HHW, the Oxford NIHR Comprehensive Biomedical Research Centre, core funding to the Wellcome Trust Centre for Human Genetics from the Wellcome Trust (090532/Z/09/Z), a Wellcome Trust Clinical Training Fellowship to MG and a Health Foundation/Academy of Medical Sciences Clinician Scientist Fellowship award to DNC. The views expressed are those of the authors and not necessarily those of the NHS, the NIHR, the Department of Health or the Wellcome Trust.

## Author Contributions

HHW, RA and MG performed TCGA and RNAseq analyses; JML and TP performed the IHC and IF staining; PV and KL performed the RNAseq experiments; TB, CLC, and IG collected, processed and selected the tumor tissue for IHC and IF; FAE, MCAW, FLK and EP collected and processed tumor tissue for RNAseq and ex vivo studies; HHW, JML and AK performed the in vitro and ex vivo analyses; DNC, HWN, and MB supervised the study; HHW, JML, HWN and MB conceived and designed the study and wrote the paper.

